# YY1 Binding Motif Upstream of Rep/Cap Increases AAV Yield and Full Capsids

**DOI:** 10.64898/2026.05.21.726733

**Authors:** Yuri Ofusa, Sanae Nishio, Tatsuji Enoki, Junichi Mineno, Keiya Ozawa, Hiroaki Mizukami, Kenji Ohba

**Affiliations:** Division of Genetic Therapeutics, Center for Molecular Medicine, Jichi Medical University, Shimotsuke, Tochigi, Japan; CDM Center, TAKARA Bio Inc., Kusatsu, Shiga, Japan; Center for Gene Therapy Research, Jichi Medical University, Shimotsuke, Tochigi, Japan

**Keywords:** Adeno-associated virus, AAV vector, Gene therapy, Replicase/Capsid, Yin Yang 1 (YY1), Vector yield, Empty/full capsid ratio, Packaging efficiency

## Abstract

Adeno-associated virus (AAV) vectors are widely used in gene therapy, whereas low manufacturing efficiency and a large proportion of empty capsids are major obstacles. This study focused on the Yin Yang 1 (YY1) binding motif (YY1-motif) and investigated the effect of its presence or insertion upstream of the Replicase (Rep)/Capsid (Cap) gene on AAV vector production. We found that the YY1-motif incidentally presented in a Rep/Cap plasmid was associated with high vector production. We then designed several modified Rep/Cap (RC2) constructs. The YY1-motif insertion in front of Rep/Cap gene increased vector yield in a repeat-number-dependent manner, and similar effects were not observed with other promoters insertion. Furthermore, the insertion of the YY1-motif reduced the amount of Cap protein per the same amount of full particle in supernatants on multiple serotypes, indicating the improvement in the empty/full capsid ratio. The YY1-motif insertion did not affect the AAV vector infectivity. These results denote that the YY1-motif has a universal regulatory function that optimizes the Rep/Cap expression balance, and simultaneously improves the production efficiency and full particle formation of AAV vectors. This finding could contribute to the development of highly efficient and high-quality AAV manufacturing processes.

## Introduction

Adeno-associated virus (AAV) vector is the most widely used as a gene therapy tool due to their long-term gene expression and high safety^1^. Currently, eight AAV-based gene therapy products have been approved by the FDA and EMA, and there are reported to be over 300 AAV-based clinical trials in the world^2,3^. Although AAV vector therapy has become feasible, high manufacturing costs remain a major obstacle to widespread use in clinical applications because of requiring the high dose of vector for sufficient therapeutic effects and the low production efficiency including the generation of empty particles on AAV production. In particular, the low yields and purification process by high rates of empty capsid contamination increase manufacturing costs further, pointing out that improving the manufacturing efficiency of AAV and increasing the full capsid ratio are urgent priorities to accelerate the social implementation of gene therapy.

The method of triple plasmids transfection (Helper, replicase/capsid (Rep/Cap), and AAV-gene of interest (GOI) plasmids) to HEK293 cells is widely used as the primary method for producing AAV vectors^4,5^. In the standard configuration of this system, the original Rep and Cap genes are provided with their native promoters (p5, p19, p40), but other cis-acting elements are removed to reduce the possibility of encapsulation^6^. Various studies attempt to increase vector yield, including scaling up AAV genes, promoters, cell lines, and inoculation rates, optimizing transfection, and optimizing culture media^7,8,11^. Since the expression levels and timing of Rep/Cap directly affect capsid formation and genome packaging efficiency, controlling them is a crucial factor in improving yield and complete capsid ratio^9–11^.

The full/empty capsid ratio (F/E ratio) varies depending on the vector production system, whereas the ratio of full particles in vectors produced by transfection is typically less than 20%^12–14^. Empty capsids reduce therapeutic efficacy due to increased immunogenicity and competition at target sites^15,16^, requiring the purification steps to eliminate them before clinical use. Since the purification process to remove empty capsids is directly related to high cost for gene therapy using AAV vectors, achieving the high yield and full capsid ratio in the AAV manufacturing process is extremely important^12^.

In this study, we attempted to establish a novel design that placing a portion of the promoter in front of the Rep/Cap gene in structural plasmid to increase AAV vector yield and reduce empty capsids.

## Results

### YY1 Binding Motif Located Upstream of the Rep/Cap Gene Increases AAV Vector Yield

The Rep/Cap plasmid widely used for AAV production differs from the wild-type AAV genome structure in that the original p5 promoter has been moved from upstream to downstream of the Rep/Cap gene. The p5 promoter contains a major late transcription factor (MLTF) site at position −80, a Yin Yang 1 (YY1) binding motif (YY1-motif) at position −60 (YY1−60), a Rep binding element (RBE) at position −20 and a YY1-motif at position +1 (YY1+1)^17^. During the analysis of various conventional Rep/Cap plasmids, we discovered a construct in which a portion of the p5 gene, YY1+1, was incidentally present in the upstream region where the p5 promoter had originally been located (Figure S1A, C). Interestingly, plasmids containing this residual sequence were found to have a higher vector yield compared to other constructs (pAAVRep2/Cap1(pAAV2/1); residual YY1 v.s. pAAV-RC1; non-YY1 upstream of Rep/Cap gene) (Figure 1A), raising a possibility that the upstream region of the Rep/Cap gene may influence AAV vector production. Therefore, we focused on this upstream region, and established a plasmid deleted YY1 (delta YY1; dYY1) (named for pAAV2/1_dYY1) from pAAV2/1 (Figure S1B) and novel Rep2/Cap2 (RC2) constructs by rearranging various promoter fragments including a portion of the p5 promoter. First, to confirm the effect of the YY1-motif-retaining pAAV2/1 plasmid, which showed high vector yield in the previous analysis, AAV vectors were produced with conventional triple plasmids transfection method using pAAV2/1 or pAAV2/1_dYY1. The deletion of residual YY1-motif in front of Rep/Cap gene reduced the vector yield compared with original pAAV2/1 (Figure 1B, C), indicating that the insertion of promoter or regulatory element upstream of Rep/Cap construct could improve AAV vector productivity. Next, a modified plasmid (pAAV-RC2_InsYY1) was created by the insertion of a YY1-motif in front of the Rep/Cap in a pAAV-RC2 plasmid (pAAV-RC2) that lacked the YY1-motif. AAV vector yield was increased by YY1-motif insertion with the opposite manner as compared to the case of YY1 deletion (Figure 1D, E). These results indicate that the YY1-motif located upstream of the Rep/Cap enhances AAV vector production even in the RC2 backbone.

**Figure 1.**
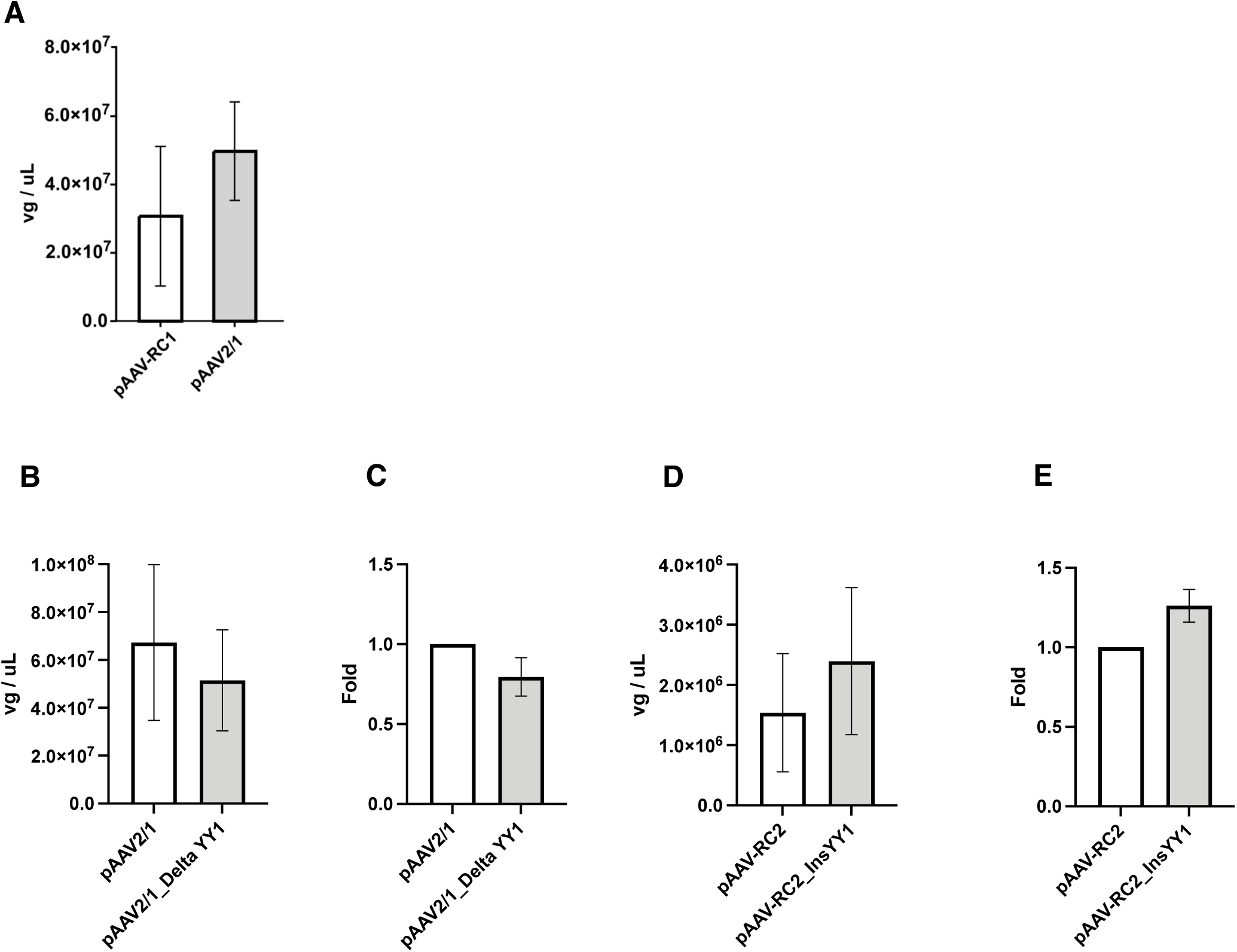
YY1 Binding Motif Located Upstream of the Rep/Cap Gene Increases AAV Vector Yield. **A-E** Adeno-associated virus (AAV) vector yield. (A, B, D) Amount of AAV vector in solution (vg/mL). vg = vector genome. (C, E) Fold differences in AAV vector yield. Data were calculated from three independent experiments and normalized with the value of the pAAV-2/1 WT(C) and pAAV-RC2 WT(E). Error bars indicate the standard deviation (SD) of the mean.

### Promoter and Regulatory Elements Located Upstream of the Rep/Cap Gene Differentially Affect AAV Vector Yield, Rep/Cap Expression, and Infectivity

Our results indicate a possibility that the insertion of promoter or regulatory element including YY1-motif upstream of Rep/Cap construct can improve the AAV productivity to some extent. We then investigated whether various promoter and regulatory elements located upstream of Rep/Cap in the RC2 could promote the production of AAV vectors (Figure 2A). When the repeat number of the YY1-motif was increased with 1, 2, or 4, the vector yield increased accordingly (Figure 2B, C). Contrarily, the addition of other promoters (CMV or CAG promoters) in front of the Rep/Cap gene showed the no increase of vector production. In addition, focusing on the fact that YY1 is located at the downstream most sequence of the AAV2 p5 promoter, we introduced the minimal promoter (miniP), which corresponds to the downstream region of CMV, and the transcription factor complex NF-κB as comparison elements. As a result, although the addition of these promoters or elements in front of the Rep/Cap gene slightly increased vector yield, especially in miniP, compared to the wild type (Wt), they did not exhibit enhancement effect with the same level as the YY1-motif. These results indicate that YY1-motif insertion upstream of Rep/Cap gene somehow improve the AAV vector production. Western blot analysis of Rep and Cap protein expression levels revealed that plasmids containing the YY1-motif showed a slight decrease and increase in Rep78 and Rep68 expression respectively compared to the reference RC2 plasmid, while Rep52 expression remained almost unchanged (Figure 2D). For Cap proteins, VP1, VP2, and VP3 all showed reduced expression levels in plasmids containing the YY1-motif with no change of VP1:VP2:VP3 ratio. On the other hand, the signals for Rep78 and Rep68 were strongly detected, while Rep52 was weak in RC2-CMV and RC2-CAG. Furthermore, all Cap proteins (VP1, VP2, VP3) showed remarkably weak signals in CMV and CAG groups. RC2-miniP showed a similar expression pattern to RC2-CMV and RC2-CAG. Additionally, Rep78 and Rep68 were slightly enhanced in RC2-4xNF-κB.

**Figure 2.**
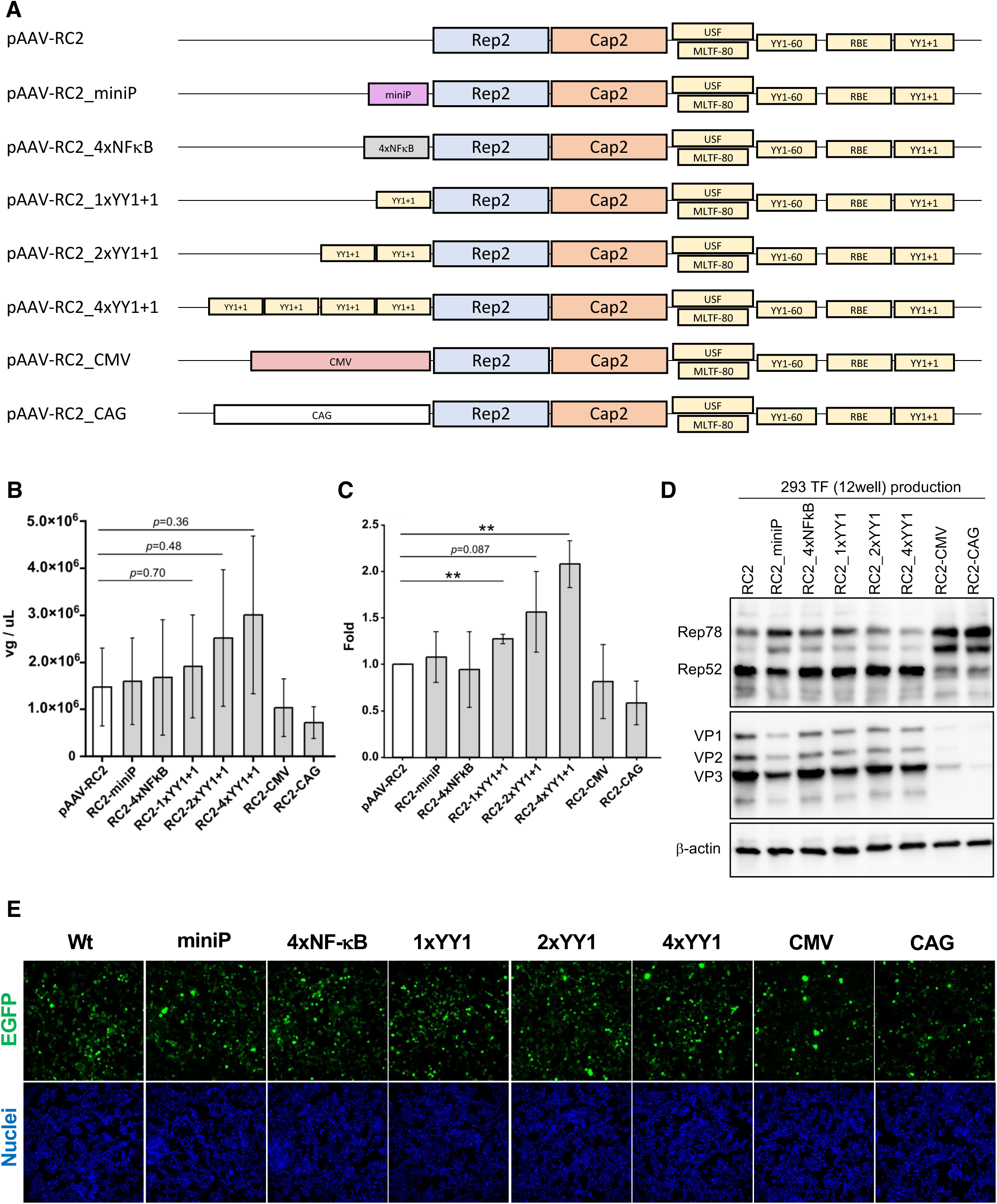
Promoter and Regulatory Elements Located Upstream of the Rep/Cap gene Differentially Affect AAV Vector Yield, Rep/Cap Expression, and Infectivity. **A** Schematic representation of modified Rep/Cap2 plasmid constructs containing various promoter motif upstream of the Rep/Cap genes. **B, C** AAV vector yield using modified Rep/Cap2 plasmid constructs. (A) Amount of AAV vector in solution (vg/mL). (B) Fold differences in AAV vector yield. Data were calculated from three independent experiments and normalized with the value of pAAV-RC2 WT. Statistical significance is indicated as: ** p < 0.01. Exact p-values are shown where applicable. Error bars indicate the standard deviation (SD) of the mean. **D** Western blot of replicase (Rep) and capsid (Cap) proteins in HEK293 cells. Rep, Cap and b-actin proteins were detected using anti-Rep, anti-Cap and anti-b-actin antibodies, respectively. **E** Infectivity data of AAV vectors in vitro. The 2v6.11 cells were infected with AAV2 (500 vg/cell). Cells were observed at 96 h after infection. Data indicate the infectivity of the AAV vector (EGFP; green) in AAV2.

Since the Cap proteins VP1 and VP2 have the same amino acid sequence as VP3, plus an additional N-terminal sequence necessary for infection^18^, changes in Cap protein levels may affect infectivity. However, when comparing the infectivity of vectors produced by each construct, similar infection efficiencies were obtained in all constructs (Figure 2E), meaning that the decrease in Cap protein levels observed in Western blot did not significantly affect infectivity, at least under these conditions.

These data denote that YY1-motif insertion upstream of Rep/Cap gene can improve the AAV vector yield without affecting viral vector function.

### YY1 Binding Motif Upstream of the Rep/Cap Gene Reduces Empty Capsid in RC2

Plasmids containing the YY1-motif showed increased viral vector yield while intracellular Cap protein levels decreased, raising a possible reduction in Cap protein per viral vector. Therefore, using 2 × 10^8^ vg of AAV vectors (quantified vg by qPCR) in the RC2 serotype, immunoprecipitation was performed using A20 antibody and protein A/G magnetic beads to know particle formation efficacy as performed in our previous work showing that the data is corresponding to the one of empty capsid level calculated by qPCR/ELISA^19^. Cap protein levels significantly decreased along with increasing YY1-motif levels (Figure 3A). Because Cap proteins are known to exist in a ratio of VP1:VP2: VP3 = 1:1:10^20^, the band intensity of VP3 was quantified in this study as an indicator of Cap protein levels (Figure 3B). These results indicate that the F/E capsid ratio, consistent with the Cap protein amount ratio obtained by Western blotting after qPCR/IP was improved, meaning that the introduction of the YY1-motif enhances full particle formation.

**Figure 3.**
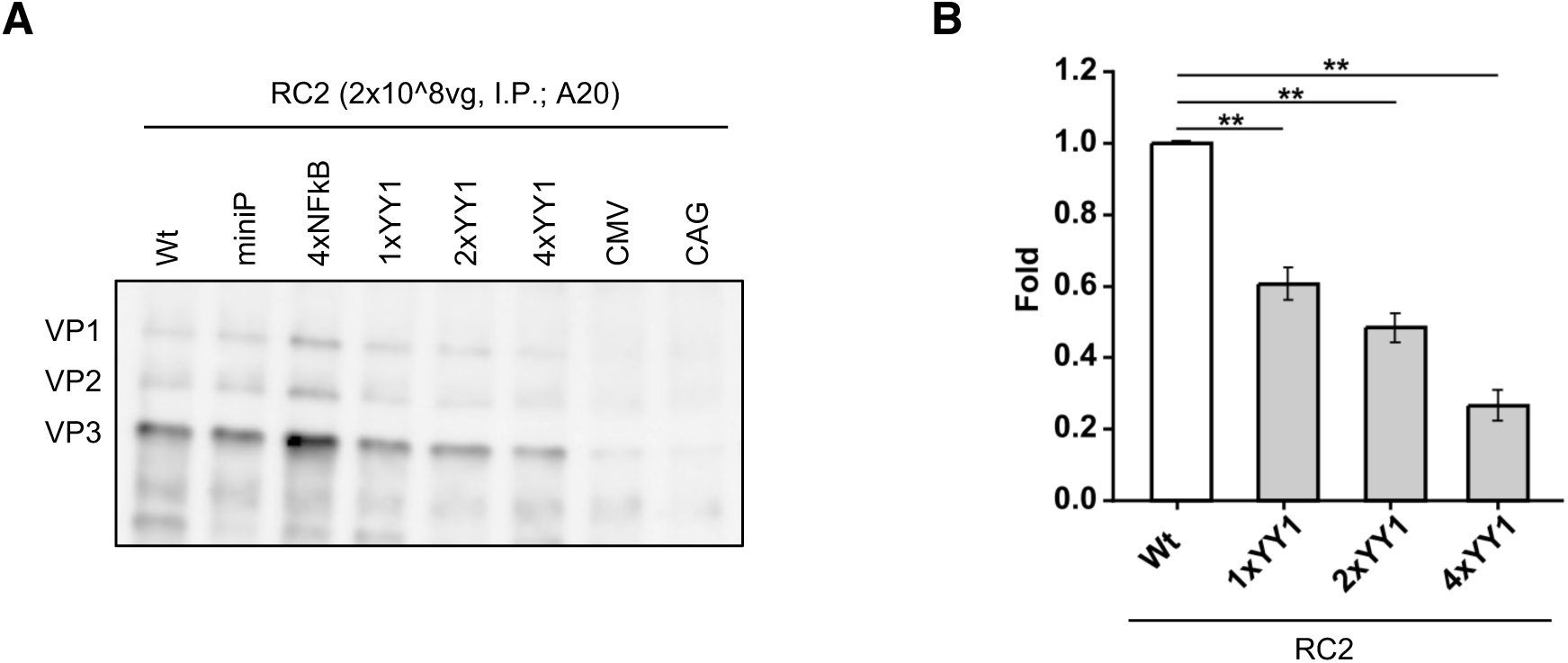
YY1 Binding Motif Upstream of the Rep/Cap Gene Reduces Empty Capsid in RC2. **A** Representative data on western blot of capsid (Cap) proteins after immunoprecipitation. The same titer of adeno-associated virus (AAV) vector (2×10^8^ vg/sample) calculated using qPCR was subjected to immunoprecipitation using A20 antibodies and protein A and G (A/G) magnetic beads before western blotting. Cap proteins were detected using anti-Cap antibodies. **B** VP3 band intensities after immunoprecipitation. Data were calculated from three independent experiments and normalized with the value of Wt. Statistical significance is indicated as: ** p < 0.01. Error bars indicate the standard deviation (SD) of the mean.

### YY1 Binding Motif Located Upstream of the Rep/Cap Gene Increases AAV Vector Yield in Multiple Serotypes

Introducing the YY1-motif into pAAV-RC2 increased AAV vector yield and improved the F/E ratio. Next, we investigated whether this phenomenon occurred in other AAV serotypes. Comparing the Wt plasmid with a plasmid containing four repeat of the YY1-motif (4×YY1), we observed an increase in vector yield in all serotypes, although the degree of increase varied (Figure 4A-F). Furthermore, similar to Figure 1B and C, removing the YY1-motif (deleted YY1; deltaYY1) from AAV2/1, which originally contained the YY1-motif upstream of the Rep/Cap gene, resulted in a decrease in vector yield (Figure 4G, H).

**Figure 4.**
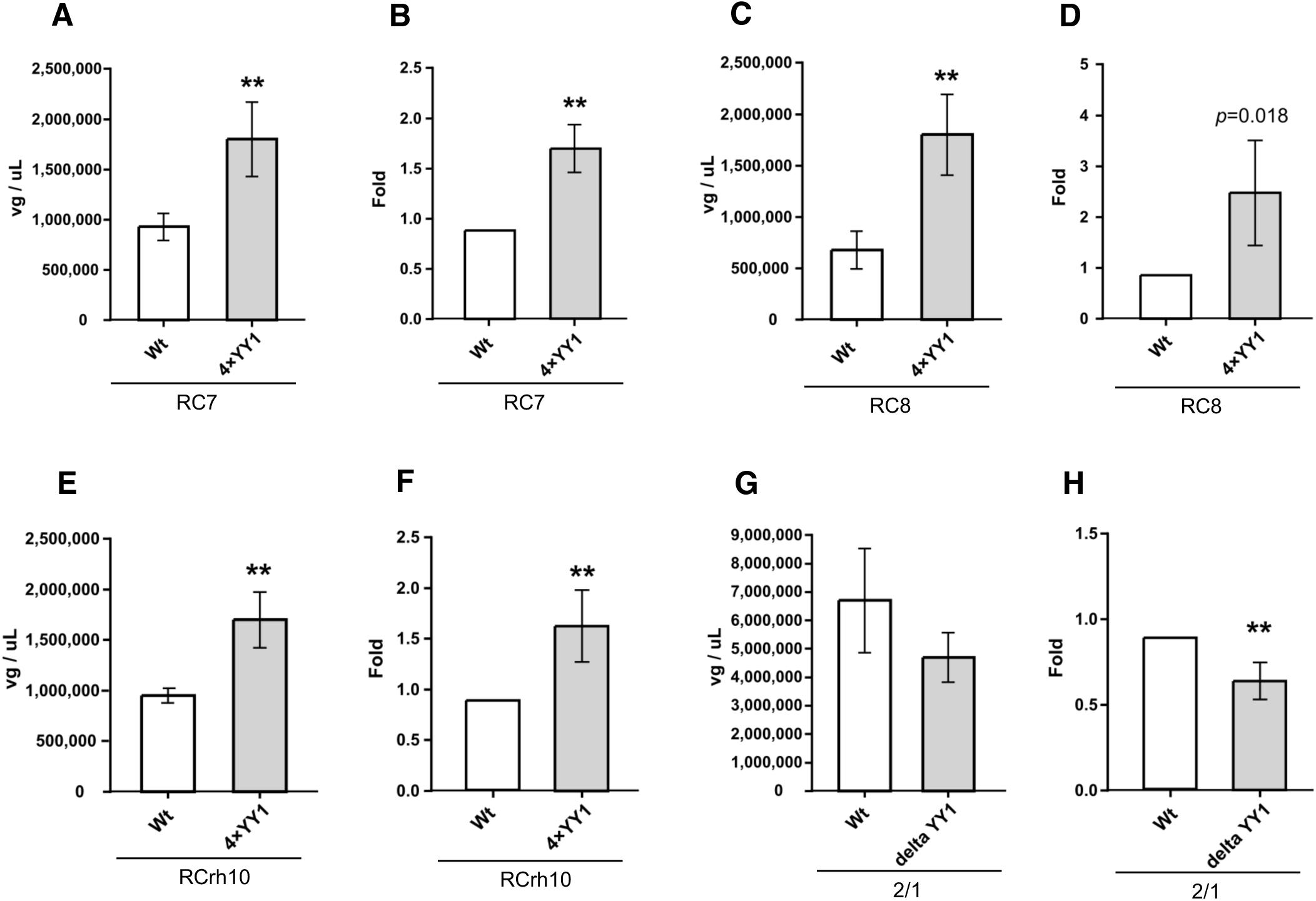
YY1 Binding Motif Located Upstream of the Rep/Cap Gene Increases AAV Vector Yield in Multiple Serotypes. **A-H** AAV vector yield using WT and 4×YY1-Containing Plasmids. (A, C, E, G) Amount of AAV vector in solution (vg/mL). (B, D, F, H) Fold differences in AAV vector yield. Data were calculated from four independent experiments and normalized with the value of each corresponding WT. Statistical significance is indicated as: ** p < 0.01. Exact p-values are shown where applicable. Error bars indicate the standard deviation (SD) of the mean.

### YY1 Binding Motif Located Upstream of the Rep/Cap Gene Reduces Empty Capsid in Multiple AAV Serotypes

Additionally, immunoprecipitation was performed using the same number of viral genomes for each serotype, and the amount of Cap protein was compared (Figure 5A-E). A decrease in Cap protein was observed in the 4×YY1 plasmid for all serotypes. Conversely, Cap protein increased in delta YY1 (dYY1). These results indicate that the F/E ratio varies depending on the presence or absence of the YY1-motif. Furthermore, regarding infectivity, similar to RC2, no changes were observed due to the introduction or removal of the YY1-motif (Figure 5F).

**Figure 5.**
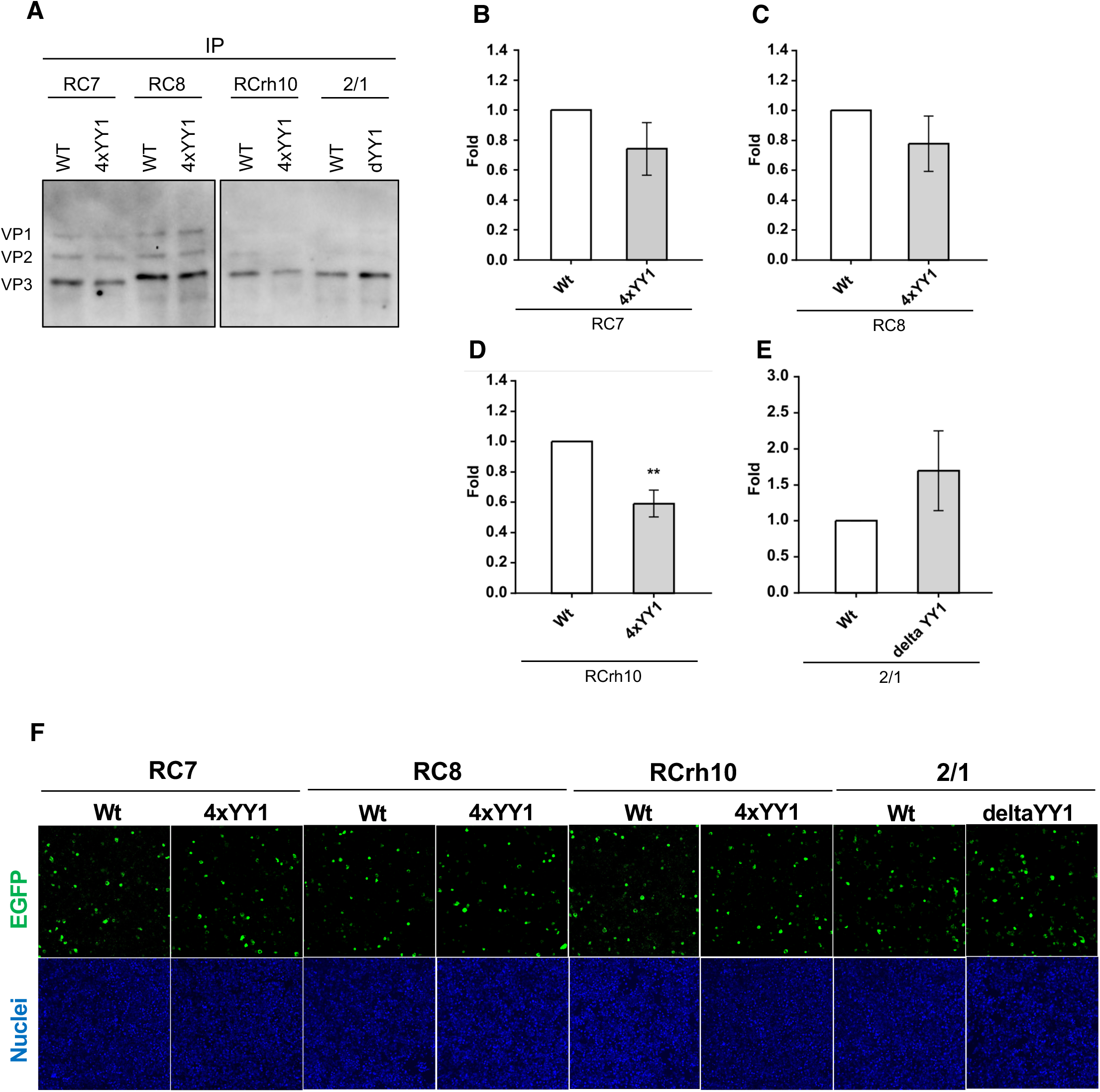
YY1 Binding Motif Located Upstream of the Rep/Cap Gene Reduces Empty Capsid in Multiple AAV Serotypes. **A** Western blot of capsid (Cap) proteins after immunoprecipitation. The same titer of adeno-associated virus (AAV) vector (5×10^8^ vg/sample) calculated using qPCR was subjected to immunoprecipitation using ADK8(RC7, RC8, RC10) and ADK1a (2/1) antibodies and protein A and G (A/G) magnetic beads before western blotting. Cap proteins were detected using anti-Cap antibodies. **B-E** VP3 band intensities after immunoprecipitation. Data were calculated from three independent experiments and normalized with the value of each corresponding WT. Error bars indicate the standard deviation (SD) of the mean. F Infectivity data of AAV vectors in vitro. The 2v6.11 cells were infected with AAV7 (785 vg/cell), AAV8 (692 vg/cell), AAVrh10 (797 vg/cell) and AAV2/1 (1010 vg/cell). Cells were observed at 72 h after infection. Data indicates the infectivity of the AAV vector (EGFP; green) in AAV2.

These results demonstrate that the YY1-motif could be a regulatory factor that broadly influences AAV vector production and full particle formation, independent of specific serotypes.

## Discussion

This study revealed that a YY1 binding motif (YY1-motif) at upstream of the Rep/Cap gene significantly increased AAV vector yield. This effect was not observed with other promoters or transcription factors, meaning that it’s a YY1-specific phenomenon. Moreover, the YY1-motif insertion is effective for increasing the yield in multiple AAV serotypes, whereas the infectivity of AAV vectors was not affected. In addition, the introduction of the YY1-motif reduced the amount of Cap protein per viral genome, suggesting that the upstream region of the Rep/Cap gene may influence genome packaging efficiency.

The decrease in vector production when strong promoters such as CMV and CAG were placed upstream of the Rep/Cap gene suggests that stronger Rep/Cap expression is not always better, and that there is an optimal expression level and timing for pAAV-Rep/Cap (Figure 2). Furthermore, while miniP and NF-κB elements increased yield more than Wt, they did not show the same effect as the YY1-motif. This suggests that YY1 may not only function as a transcriptional activation but also work in coordination with the p5 promoter and subsequent p19 activation. Furthermore, the increase in vector yield with increasing repeat number of the YY1-motif implies that YY1 contributes directly and depends on the amount of recruited YY1 to target regions to the regulation of Rep/Cap gene expression. Present study also revealed that changes in the expression balance of Rep/Cap proteins due to differences in promoters and regulatory sequences are related to vector production efficiency (Figure 2). The two large Rep proteins (Rep78/68) are promoted by promoter p5 and are involved in DNA replication and regulation of the viral promoter^18,21^. The two small Rep proteins (Rep52/40) are activated by the p19 promoter activated by Rep78/68 and are essential for genome accumulation and packaging of DNA into the capsid^22^. Cap proteins (VP1, VP2, VP3) are initiated by the p40 promoter^2^. In this study, strong promoters such as CMV and CAG showed overexpression of Rep78/68 and a significant decrease in Cap expression (Figure 2D). This result is consistent with previous reports that overexpression of Rep78/68 inhibits AAV DNA replication and cap gene expression, and conversely, that reducing Rep78/68 levels increases vector genome titer^23,24^. On the other hand, under conditions where the YY1-motif was introduced, Rep78 decreased and Rep68 slightly increased while Rep52 was maintained, and Cap protein levels also decreased overall.

Furthermore, the fact that miniP showed a pattern similar to CMV/CAG, and that Rep78/68 was slightly enhanced with the NF-κB sequence, is consistent with the reason why the vector yield did not increase as much under these conditions as with the YY1-motif. In addition, some reports show that improving the Rep/Cap plasmid involves introducing the p5 promoter downstream of Rep/Cap, which slightly weakens Rep78 expression while maintaining the expression of p19 and p40 and maximizing their titers^6^. In this study as well, it is thought that inserting YY1, which is part of p5, upstream of Rep/Cap while p5 is present downstream, moderately suppressed Rep78 while maintaining Rep52, which contributed to the increase in yield. The expression levels and balance of Rep/Cap are immensely important in AAV vector production, and this construct, which inserts the YY1-motif upstream of Rep/Cap, may contribute to its optimization.

While the promoters and regulatory elements in front of the Rep/Cap gene altered the Rep/Cap expression pattern and vector yield, no change was observed in infectivity (Figure 2E, 5F), indicating that even though Cap proteins were reduced by the addition of the YY1-motif, infection efficiency, such as receptor binding and cell entry, was not affected.

Additionally, we revealed that when plasmids containing the YY1-motif were used, the amount of Cap protein per vg decreased stepwise with increasing YY1 levels, meaning that the introduction of the YY1-motif may contribute to an improvement in the F/E ratio, in other words, an increase in the proportion of mature particles containing AAV genomes (Figure 3, 5A-E). In particular, the fact of a promoted full particle formation even under condition of relatively low levels of intracellular Cap protein is an important finding, pointing out that the YY1-motif may be involved in increasing packaging efficiency.

The insertion of the YY1-motif increased vector yield and improved the F/E ratio in multiple serotypes (Figure 2-5). The infectivity was no change with either the introduction or removal of the YY1-motif. These results reflect that the YY1-motif may be a serotype-independent regulator. Therefore, the insertion of the YY1-motif can optimize the AAV manufacturing process by improving vector productivity and full particle formation while maintaining infectivity.

YY1 is a known transcription factor expressed throughout mammalian cells that has been reported to affect promoter activity and chromatin structure^25,26^. The molecular mechanism by which the YY1-motif increases AAV vector production and enhances packaging efficiency is unclear yet. As possible mechanisms, firstly, it can be thought that the yield was increased by mild suppression of the Rep78 protein because a previous report has shown that the YY1 protein interacts with the p5 promoter at two sites and partially suppresses p5 activity^27^. Secondly, it may be a reason that YY1 promoted p19 activation and increased packaging efficiency, since mutational analysis of the p5 promoter has demonstrated that not only YY1-60 but also YY1+1 may be essential for p19 activation^17^. However, although there is no direct evidence yet, further analyses are required. Taken together, the YY1 binding motif upstream of the Rep/Cap gene is a universal regulatory element that simultaneously improves vector yield and F/E ratio in multiple AAV serotypes, and a feasible factor to contribute to the efficiency and quality improvement of AAV vector production. Further investigation about the molecular mechanisms how YY1 influences Rep/Cap expression and packaging is expected to lead to the development of further AAV vector production systems.

## Materials and Methods

### Cell culture

HEK293 cells (Cat#RCB1637;RRID:CVCL_0045) were obtained from the Cell Engineering Division (RIKEN cell bank, Tsukuba, Japan). HEK293 and 2v6.11 cells^28^ were cultured in Eagle’s minimum essential medium (FUJIFILM Wako, Osaka, Japan) supplemented with 10% fetal bovine serum (ThermoFisher Scientific, Waltham, MA),1× MEM non-essential amino acids (FUJIFILM Wako), and 1×penicillin-streptomycin (FUJIFILM Wako).

### Plasmid construction

The pAAV2/1 (Addgene plasmid # 112862 ; http://n2t.net/addgene:112862 ; RRID:Addgene_112862), pAAV2/7 (Addgene plasmid # 112863 ; http://n2t.net/addgene:112863 ; RRID:Addgene_112863), pAAV2/8 (Addgene plasmid # 112864 ; http://n2t.net/addgene:112864 ; RRID:Addgene_112864), pAAV2/9n (Addgene plasmid # 112865 ; http://n2t.net/addgene:112865 ; RRID:Addgene_112865) and pAAV2/rh10 (Addgene plasmid # 112866 ; http://n2t.net/addgene:112866 ; RRID:Addgene_112866) for AAV serotypes 1, 7, 8, 9 and rh10 capsid (Cap) were gifted by James M. Wilson. To adjust the plasmid backbone, the Cap gene of serotype 2 in pAAV-RC2 (Agilent, Santa Clara, CA) was changed to a Cap gene from each serotype mentioned above and named pAAV-RC1, 6, 7, 8, and rh10, respectively.

For the reporter constructs, EGFP-P2A-NanoLuc (NLuc) genes (Promega, WI) were cloned into pAAV-MCS (Agilent) and named pAAV-EGFP-P2A-Nluc.

### Plasmid transfection and AAV vector production

HEK293 cells were seeded at concentrations of 8×10^5^ or 3.5×10^6^ cells/well into 24- or 6- well plates, respectively, one day before transfection. Under normal conditions, 0.5 µg pAAV-EGFP-P2A-Nluc, 0.5 µg pHelper (Cat# 6650, Takara Bio, Shiga, Japan), and 0.5 mg pAAV-RC were co-transfected into cells by using HilyMax transfection reagent (DOJINDO, Kumamoto, Japan) in 24-well plates according to the manufacturer’s instructions. All conditions were the same for co-transfection in 6-well plates except 2 µg all other components were used. For both conditions, media were replaced with fresh medium 9h after transfection. At 72h after medium change, cells and supernatants were collected in tubes. To harvest the AAVvectors, collected cells and supernatants were frozen and thawed four times, and the supernatants collected as AAV vector solutions after centrifugation and filtration by a 0.45 μm filter.

### AAV vector genome detection and titration using qPCR

For AAV vector genome detection in cell lysates, collected samples were directly applied for qPCR analysis to monitor the total replicated AAV genomic DNA in cells during AAV vector production. For AAV vector titration, collected samples were treated with TURBO DNase (Ca#AM2238, ThermoFisher Scientific) according to the manufacturer’s instructions before qPCR analysis. The total AAV vector genome and titer were quantified using qPCR with an inverted terminal repeat-specific primer set (50-GGAACCCCTAGTGATGGAGTT-30,50-CGGCCTCAGTGAGCGA30)^29^ and PowerTrack™ SYBR Green Master Mix for qPCR (Cat#A46109, Thermo Fisher Scientific) according to the manufacturer’s instructions.

### Western blotting

Cells were collected and lysed using cell lysis buffer (50 mM Tris-HCl at pH 7.5, 150 mM NaCl, and 1% NP-40). One volume of 2×sodium dodecyl sulfate (SDS) sample buffer (125 mM Tris-HCl at pH 6.8, 4% SDS, 20% glycerol,10% 2-mercaptoethanol, and 0.01% bromophenol blue) was added to the lysed samples and boiled for 10 min. Samples were subjected to a 4–20% gradient TGX precast gel (Bio-Rad Laboratories, Hercules, CA), followed by the transfer of proteins to Immobilon-P polyvinylidene fluoride membranes (Millipore, Burlington, MA) using the Trans-Blot Turbo Transfer System (Bio-Rad). After blocking membranes with 5, skim milk/TBS-T(50mMTris-HCl at pH7.6, 150mM NaCl, and 0.05% Tween20), primary antibodies were reacted at RT for 1 h, followed by secondary antibody reactions at RT for 1 h. Proteins were detected using a Clarity Western ECL substrate (Bio-Rad Laboratories) and imaged using a ChemiDoc Toch MP Imaging System (RRID:SCR_019037, Bio-Rad Laboratories). The band intensity was analyzed using ImageJ software. The following antibodies were utilized: AAVVP1/VP2/VP3rabbitpolyclonal,VP51(Cat#61084, PROGEN, Heidelberg, Germany), AAV2Replicasemousemonoclonal,303.9(Cat#61069, PROGEN), b-Actin(13E5) Rabbit(Cat#4970;RRID:AB_2223172, Cell Signaling Technology Inc, Danvers, MA), Anti-mouse IgG, HRP-linked Antibody (Cat#7076;RRID:AB_330924, Cell Signaling Technology), and Anti-rabbit IgG, HRP-linked Antibody (Cat#7074;RRID:AB_2099233, Cell Signaling Technology).

### Immunoprecipitation of AAV particles

For calculation of the full/empty particle ratio, AAV1 intact particle antibody, ADK1a (Cat#610150, PROGEN), AAV2 intact particle antibody, A20 (Cat#65155, PROGEN), and AAV8 intact particle antibody, ADK8 (Cat#610160); PROGEN) were mixed with protein A and G (A/G) magnetic beads (Cat#10001D and Cat#10003D, Dynabeads; ThermoFisher Scientific) in immunoprecipitation buffer (50 mM Tris-HCl at pH 7.5, 150 mM NaCl, and 0.05% Tween 20) at RT for 1 h. Equal amounts of AAV vectors from qPCR data (5×10^8^ vg/sample) were reacted with an antibody protein A/G magnetic beads mixture at RT for 1h with rotation. After the reaction, samples were placed in a magnetic stand to discard the buffer, and each sample washed with fresh immunoprecipitation buffer. After three times washes, pellets were boiled in 1×SDS sample buffer diluted with cell lysis buffer for 10 min, followed by western blotting. AAV capsids were detected using anti-AAV VP1/VP2/VP3 rabbit polyclonal VP51(Cat#61084, PROGEN) and anti-rabbit IgG-HRP (Cat#7074;RRID:AB_2099233, Cell Signaling technologies Inc) antibodies.

### AAV vector infection

AAV vectors were added to 2v6.11 cells with the same titer of viruses calculated from the qPCR data in 96-well plates. At 3–5 days post-infection, the infectivity of AAV vectors carrying the EGFP reporter was observed with fluorescent or confocal microscopes (Single Cellome™ System SS2000; Yokogawa Electric Corporation, Tokyo, Japan) after nuclear staining with Hoechst 33342 (ThermoFisher Scientific).

### Quantification and statistical analysis

All in vitro experiments were conducted at least three times under similar conditions.

Graphs and statistical analyses were performed using GraphPad Prism v8 software (GraphPad, San Diego, CA) and R (version 4.5.2). Means of two groups were compared using the t-test with Welch’s correction. Error bars represent standard deviation of the mean. Statistical significance is indicated in figures as follows: ** p< 0.01.

### Data Availability Statement

The authors declare that the data supporting the findings of this study are available in the article. All relevant data are available from the authors upon reasonable request.

## Acknowledgements

We thank all of our laboratory members, especially Akemi Takada, who supported the handling of our budgets. This work was supported by AMED under Grant Number, JP23bm132300, KAKENHI Grant-in-Aid for Scientific Research (B) and (C) Grant Number 26K02298 and 20K07681, and partly funded by a Joint Research Fund sponsored by Takara Bio Inc.

## Author Contributions

K.Ohba conceived and designed the study. Y.O., S.N. and K.Ohba performed the experiments and analyzed the data. Y.O. and K.Ohba interpreted experimental results. Y.O. and K.Ohba wrote the manuscript. Y.O., T.E., J.M., K. Ozawa, H.M. and K.Ohba edited the manuscript. All the authors have reviewed and approved the final manuscript.

## References

1. Wang, J.-H., Gessler, D.J., Zhan, W., Gallagher, T.L., and Gao, G. (2024). Adeno-associated virus as a delivery vector for gene therapy of human diseases. Sig Transduct Target Ther 9, 78. 10.1038/s41392-024-01780-w.

2. Suarez-Amaran, L., Song, L., Tretiakova, A.P., Mikhail, S.A., and Samulski, R.J. (2025). AAV vector development, back to the future. Molecular Therapy 33, 1903–1936. 10.1016/j.ymthe.2025.03.064.

3. ClinicalTrials.gov U.S. National Library of Medicine. https://clinicaltrials.gov/.

4. Grimm, D., Kern, A., Rittner, K., and Kleinschmidt, J.A. (1998). Novel Tools for Production and Purification of Recombinant Adenoassociated Virus Vectors. Human Gene Therapy 9, 2745–2760. 10.1089/hum.1998.9.18-2745.

5. Xiao, X., Li, J., and Samulski, R.J. (1998). Production of High-Titer Recombinant Adeno-Associated Virus Vectors in the Absence of Helper Adenovirus. J Virol 72, 2224–2232. 10.1128/JVI.72.3.2224-2232.1998.

6. Aponte-Ubillus, J.J., Barajas, D., Peltier, J., Bardliving, C., Shamlou, P., and Gold, D. (2018). Molecular design for recombinant adeno-associated virus (rAAV) vector production. Appl Microbiol Biotechnol 102, 1045–1054. 10.1007/s00253-017-8670-1.

7. Kowshik, N.C.S.S., and Singh, P. (2025). Advancing AAV vector manufacturing: challenges, innovations, and future directions for gene therapy. Front. Mol. Med. 5. 10.3389/fmmed.2025.1709095.

8. Pistek, M., Andorfer, P., Grabherr, R., Kraus, B., and Hernandez Bort, J.A. (2024). Factors affecting rAAV titers during triple-plasmid transient transfection in HEK-293 cells. Biotechnol Lett 46, 945–959. 10.1007/s10529-024-03520-0.

9. Chen, Q., Lee, C.H., Whitfield, R., and Nesbeth, D.N. (2025). Chronologically distributed transfection improves AAV2 and AAV2/8 capsid filling and reveals assembly schedule divergence. Molecular Therapy Methods & Clinical Development 33, 101610. 10.1016/j.omtm.2025.101610.

10. Emmerling, V.V., Pegel, A., Milian, E.G., Venereo-Sanchez, A., Kunz, M., Wegele, J., Kamen, A.A., Kochanek, S., and Hoerer, M. (2016). Rational plasmid design and bioprocess optimization to enhance recombinant adeno-associated virus (AAV) productivity in mammalian cells. Biotechnology Journal 11, 290–297. 10.1002/biot.201500176.

11. Lee, M., and Park, J.E. (2026). The evolving landscape of recombinant adeno-associated virus (rAAV) manufacturing: Engineering vector plasmids for upstream production. Biotechnology Advances 90, 108906. 10.1016/j.biotechadv.2026.108906.

12. Schnödt, M., and Büning, H. (2017). Improving the Quality of Adeno-Associated Viral Vector Preparations: The Challenge of Product-Related Impurities. Human Gene Therapy Methods 28, 101–108. 10.1089/hgtb.2016.188.

13. Allay, J.A., Sleep, S., Long, S., Tillman, D.M., Clark, R., Carney, G., Fagone, P., McIntosh, J.H., Nienhuis, A.W., Davidoff, A.M., et al. (2011). Good Manufacturing Practice Production of Self-Complementary Serotype 8 Adeno-Associated Viral Vector for a Hemophilia B Clinical Trial. Human Gene Therapy 22, 595–604. 10.1089/hum.2010.202.

14. Penaud-Budloo, M., François, A., Clément, N., and Ayuso, E. (2018). Pharmacology of Recombinant Adeno-associated Virus Production. Molecular Therapy - Methods & Clinical Development 8, 166–180. 10.1016/j.omtm.2018.01.002.

15. Mingozzi, F., Maus, M.V., Hui, D.J., Sabatino, D.E., Murphy, S.L., Rasko, J.E.J., Ragni, M.V., Manno, C.S., Sommer, J., Jiang, H., et al. (2007). CD8+ T-cell responses to adeno-associated virus capsid in humans. Nat Med 13, 419–422. 10.1038/nm1549.

16. Gao, K., Li, M., Zhong, L., Su, Q., Li, J., Li, S., He, R., Zhang, Y., Hendricks, G., Wang, J., et al. (2014). Empty virions in AAV8 vector preparations reduce transduction efficiency and may cause total viral particle dose-limiting side effects. Molecular Therapy - Methods & Clinical Development 1, 9. 10.1038/mtm.2013.9.

17. Lackner, D.F., and Muzyczka, N. (2002). Studies of the Mechanism of Transactivation of the Adeno-Associated Virus p19 Promoter by Rep Protein. Journal of Virology 76, 8225–8235. 10.1128/jvi.76.16.8225-8235.2002.

18. Samulski, R.J., and Muzyczka, N. (2014). AAV-Mediated Gene Therapy for Research and Therapeutic Purposes. Annu. Rev. Virol. 1, 427–451. 10.1146/annurev-virology-031413-085355.

19. Ohba, K., Sehara, Y., Enoki, T., Mineno, J., Ozawa, K., and Mizukami, H. (2023). Adeno-associated virus vector system controlling capsid expression improves viral quantity and quality. iScience 26, 106487. 10.1016/j.isci.2023.106487.

20. Wang, D., Tai, P.W.L., and Gao, G. (2019). Adeno-associated virus vector as a platform for gene therapy delivery. Nat Rev Drug Discov 18, 358–378. 10.1038/s41573-019-0012-9.

21. Maurer, A.C., and Weitzman, M.D. (2020). Adeno-Associated Virus Genome Interactions Important for Vector Production and Transduction. Human Gene Therapy 31, 499–511. 10.1089/hum.2020.069.

22. King, J.A. (2001). DNA helicase-mediated packaging of adeno-associated virus type 2 genomes into preformed capsids. The EMBO Journal 20, 3282–3291. 10.1093/emboj/20.12.3282.

23. Li, J., Samulski, R.J., and Xiao, X. (1997). Role for highly regulated rep gene expression in adeno-associated virus vector production. J Virol 71, 5236–5243. 10.1128/jvi.71.7.5236-5243.1997.

24. Ogasawara, Y., Urabe, M., and Ozawa, K. (1998). The Use of Heterologous Promoters for Adeno-Associated Virus (AAV) Protein Expression in AAV Vector Production. MICROBIOLOGY and IMMUNOLOGY 42, 177–185.

25. Weintraub, A.S., Li, C.H., Zamudio, A.V., Sigova, A.A., Hannett, N.M., Day, D.S., Abraham, B.J., Cohen, M.A., Nabet, B., Buckley, D.L., et al. (2017). YY1 Is a Structural Regulator of Enhancer-Promoter Loops. Cell 171, 1573–1588.e28. 10.1016/j.cell.2017.11.008.

26. Verheul, T.C.J., van Hijfte, L., Perenthaler, E., and Barakat, T.S. (2020). The Why of YY1: Mechanisms of Transcriptional Regulation by Yin Yang 1. Front. Cell Dev. Biol. 8. 10.3389/fcell.2020.592164.

27. Pereira, D.J., McCarty, D.M., and Muzyczka, N. (1997). The adeno-associated virus (AAV) Rep protein acts as both a repressor and an activator to regulate AAV transcription during a productive infection. J Virol 71, 1079–1088. 10.1128/jvi.71.2.1079-1088.1997.

28. Mohammadi, E.S., Ketner, E.A., Johns, D.C., and Ketner, G. (2004). Expression of the adenovirus E4 34k oncoprotein inhibits repair of double strand breaks in the cellular genome of a 293-based inducible cell line. Nucleic Acids Res 32, 2652–2659. 10.1093/nar/gkh593.

29. Aurnhammer, C., Haase, M., Muether, N., Hausl, M., Rauschhuber, C., Huber, I., Nitschko, H., Busch, U., Sing, A., Ehrhardt, A., et al. (2012). Universal Real-Time PCR for the Detection and Quantification of Adeno-Associated Virus Serotype 2-Derived Inverted Terminal Repeat Sequences. Human Gene Therapy Methods 23, 18–28. 10.1089/hgtb.2011.034.

