## Supplemental Figure S1 for "YY1 Binding Motif Upstream of Rep/Cap Increases AAV Yield and Full Capsids"

**A pAAV2/1 plasmid structure**

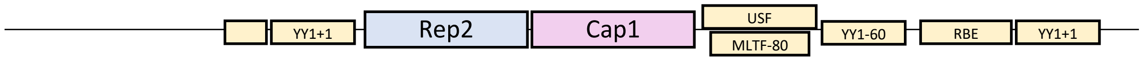


**B pAAV2/1_delta YY1 (dYY1) plasmid structure**

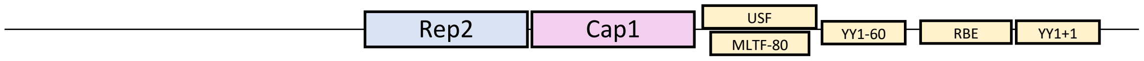


**C pAAV-RC2 plasmid structure**

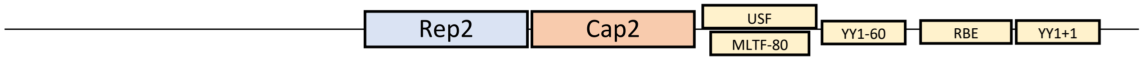


**Supplemental Figure S1. The structural difference of Rep/Cap plasmid between pAAV2/1 and pAAV-RC2.**

**A** Plasmid structure of pAAV2/1. Partial AAV promoter region remains in front of Rep2/Cap1 (Rep/Cap1) gene.
**B** Plasmid structure of pAAV2/1_dYY1**.** pAAV2/1 deletion mutant about remaining region upstream of Rep/Cap1 gene.
**C** Plasmid structure of pAAV-RC2. The common structure of pAAV- RC (Rep/Cap) plasmids including AAV2, which are commercialized widely.
